# Coevolutionary dynamics via adaptive feedback in collective-risk social dilemma game

**DOI:** 10.1101/2022.12.19.520980

**Authors:** Linjie Liu, Xiaojie Chen, Attila Szolnoki

**Affiliations:** College of Science, Northwest A & F University, Yangling 712100, China; School of Mathematical Sciences,University of Electronic Science and Technology of China, Chengdu 611731, China; Institute of Technical Physics and Materials Science, Centre for Energy Research, P.O. Box 49, H-1525 Budapest, Hungary

## Abstract

Human society and natural environment form a complex giant ecosystem, where human activities not only lead to the change of environmental states, but also react to them. By using collective-risk social dilemma game, some studies have already revealed that individual contributions and the risk of future losses are inextricably linked. These works, however, often use an idealistic assumption that the risk is constant and not affected by individual behaviors. We here develop a coevolutionary game approach that captures the coupled dynamics of cooperation and risk. In particular, the level of contributions in a population affects the state of risk, while the risk in turn influences individuals’ behavioral decision-making. Importantly, we explore two representative feedback forms describing the possible effect of strategy on risk, namely, linear and exponential feedbacks. We find that cooperation can be maintained in the population by keeping at a certain fraction or forming an evolutionary oscillation with risk, independently of the feedback type. However, such evolutionary outcome depends on the initial state. Taken together, a two-way coupling between collective actions and risk is essential to avoid the tragedy of the commons. More importantly, a critical starting portion of cooperators and risk level is what we really need for guiding the evolution toward a desired direction.

## Introduction

Human activities constantly affect the natural environment and cause changes in its quality, which in turn affects our daily life and health conditions (***Patz et al., 2005; Steffen et al., 2006***; ***Perc et al., 2017***; ***Obradovich et al., 2018***; ***Hilbe et al., 2018***; ***Su et al., 2019, 2022***). A well-known example is climate change, which is one of the biggest contemporary challenges of our civilization (***Parmesan and Yohe, 2003***; ***Stone et al., 2013***). A large number of carbon emissions caused by human activities will exacerbate the greenhouse effect, which risks raising global temperatures to dangerous levels. The direct consequences of global warming are the melting of glaciers and the rise of sea level, which will inevitably affect human activities (***Schuur et al., 2015***; ***Obradovich and Rahwan, 2019***; ***Moore et al., 2022***). Similarly, we can give more examples of coupled human and natural systems to continue this list, such as habitat destruction and the spread of infectious diseases (***Liu et al., 2001, 2007***; ***Chen and Fu, 2019***; ***Tanimoto, 2021***; ***Chen and Fu, 2022***). At present, the importance of developing a new comprehensive framework to study the coupling between human behavior and the environment has been recognized by number of interdisciplinary approaches (***Weitz et al., 2016***; ***Chen and Szolnoki, 2018***; ***Tilman et al., 2020***).

Evolutionary game theory provides a powerful theoretical framework for studying the coupled dynamics of human and natural systems (***Maynard Smith, 1982***; ***Weibull, 1997***; ***Stewart and Plotkin, 2014***; ***Radzvilavicius et al., 2019***; ***Park et al., 2020***; ***Niehus et al., 2021***; ***Han et al., 2021***; ***Cooper et al., 2021***). Furthermore, coevolutionary game models have recognized the fact that individual payoff values are closely related to the state of the environment (***Weitz et al., 2016***; ***Szolnoki and Chen, 2018***; ***Chen and Szolnoki, 2018***; ***Hauert et al., 2019***; ***Tilman et al., 2020***; ***Wang and Fu, 2020***; ***Yan et al., 2021***). For example, Weitz et al. (***Weitz et al., 2016***) considered the dynamical changes of the environment, which modulates the payoffs of individuals. Their results show that individual strategies and the environmental state may form a sustained cycle where strategy swing between full cooperation and full defection, while the environment state oscillates between the replete state and the depleted state. Along this line, feedback-evolving game systems with intrinsic resource dynamics (***Tilman et al., 2020***), asymmetric interactions in heterogeneous environments (***Hauert et al., 2019***), and time-delay effect (***Yan et al., 2021***) have been also investigated where periodic oscillation of strategy and environment are observed. As a general conclusion, the feedback loop between individual strategies and related environment is a key element to maintain long-term cooperation and sustainable use of resources.

Despite of the mentioned efforts, the research on possible consequences of the feedback between human activity and natural systems is still in early stage. Staying at the above mentioned example, potential feedback loops between human activities and climate change exist (***Obradovich and Rahwan, 2019***). However, most scholars study these two topics, that is, human contributions to climate change and social impacts of the changing climate on human behavior, in a separated way (***Vitousek et al., 1997***; ***Barfuss et al., 2020***). On the one hand, some of them usually focus on how human behaviors (use of land, oceans, fossil fuel, freshwater, etc.) affect environment (***Vitousek et al., 1997***). On the other hand, researchers who are interested in society and biology frequently focus on how environmental change will affect human behaviors (***Culler et al., 2015***; ***Obradovich and Rahwan, 2019***; ***Celik, 2020***). Recently, these two approaches have been merged into a single framework, called as collective-risk social dilemma game, which serves as a general paradigm for studying climate change dilemmas (***Milinski et al., 2008***). Within it, a group of individuals decide whether to contribute to reach a collective goal. If the total contributions of all individuals exceed a certain threshold, then the disaster is averted and all individuals benefit from it. Otherwise, the disaster occurs with a probability (also known as the risk of collective failure), resulting in fatal economic losses for all participants. Both behavioral experiments and theoretical works show that the risk of future losses plays an important role in the evolution of cooperation (***Milinski et al., 2008***; ***Santos and Pacheco, 2011***; ***Chen et al., 2012a***; ***Vasconcelos et al., 2013***; ***Hilbe et al., 2013***; ***Vasconcelos et al., 2014***; ***Barfuss et al., 2020***; ***Domingos et al., 2020***; ***Sun et al., 2021***; ***Chica et al., 2022***).

Previous studies based on the collective-risk dilemma game revealed that the risk of collective failure could affect individuals’ motivation to cooperate when they face to the problem of collective action, but ignored an important practical aspect. That is, human decision-making is not only affected by changes in the risk state, but also affects the level of risk (***Chen et al., 2012a***). Indeed, the risk of collective failure is lower in a highly cooperative society, but becomes significant in the opposite case. This fact is not only reflected in climate change (***Moore et al., 2022***), but also in the spread of infectious diseases (***Chen and Fu, 2022***) and vaccination (***Nichol et al., 1998***; ***Chen and Fu, 2019***). Furthermore, although the risk level varies in a changing population, their relation is not necessarily straightforward. For example, a study revealed that the infection-fatality risk (IFR) of COVID-19 in India decreased linearly from June 2020 to September 2020 due to improved healthcare or increased vaccination (***Yang and Shaman, 2022***). Throughout the whole process (from March 2020 to April 2021), the statistical curve of IFR is nonlinear, that is, when the epidemic broke out, the value of IFR remained at a high level, and then with the increase of vaccination or the improvement of healthcare, the IFR value gradually decreased, then flattened and remained at a low level (***Yang and Shaman, 2022***). On the other hand, the change of risk is bound to affect individuals’ decision-making, which has been confirmed in behavioral experiments and theoretical research (***Milinski et al., 2008***; ***Pacheco et al., 2014***). Though potential feedback loops between strategy and risk of future losses are already recognized, a study focusing on their direct interaction is missing. Furthermore, it is still an open question whether the character of feedback mechanism plays an essential role in the final evolutionary outcome. Hence, how the impacts of risk on human systems might, in turn, alter the future trajectories of human decision-making remains largely unexplored.

To fill this gap, we propose a coupled coevolutionary game framework based on the collective-risk dilemma to describe reciprocal interactions and feedbacks between decision-making procedure of individuals and risk. In particular, we assume that the increasing free-riding behaviors will slowly increase the risk of collective failure, and the resulting high-risk level will in turn stimulate individual contributions. However, the increase in contribution will gradually reduce the risk of collective failure, and the resulting low-risk level will promote the prevalence of free-riding behaviors again. This general feedback loop is illustrated in Fig. 1. Importantly, we respectively consider two conceptually different feedback protocols describing the effect of strategy on risk. Namely, both linear and highly nonlinear (exponential) feedback forms are checked. Our analysis identifies the conditions for the existence of stable interior equilibrium and stable limit cycle dynamics in both cases.

**Figure 1.**
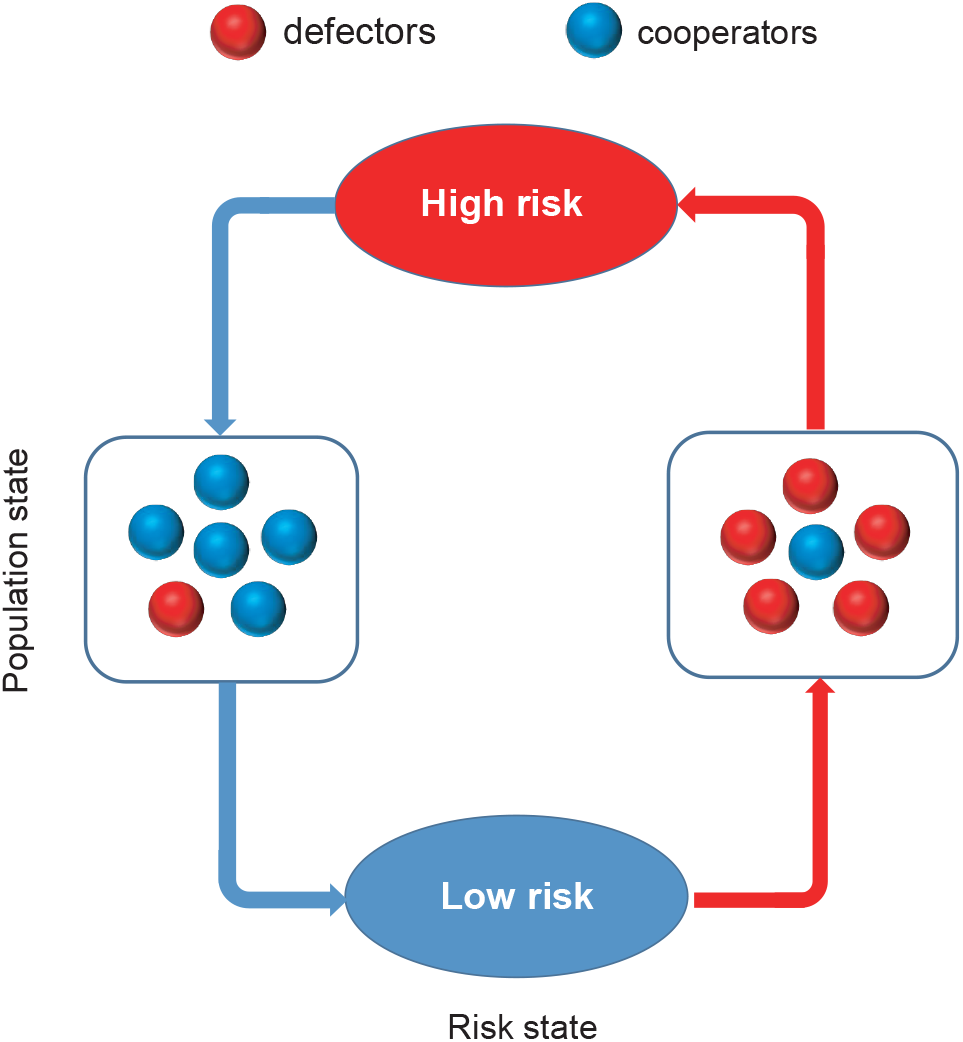
Coevolutionary feedback loop of population and risk states in the coupled game system. The meaning of colors is explained in the legend on the top.

## Methods and Materials

### Collective-risk social dilemma game

We consider an infinite well-mixed population in which *N* individuals are selected randomly to form a group for playing the collective-risk social dilemma game. Each individual in the group has an initial endowment *b* and can choose one of the two strategies, i.e., cooperation and defection. Cooperators will contribute an amount *c* to the common pool, whereas defectors contribute nothing. The remaining endowments of all individuals can be preserved if the overall number of cooperators exceeds a threshold value *M*, where 1 < *M < N* (***Milinski et al., 2008***; ***Santos and Pacheco, 2011***). Otherwise, individuals will lose all their endowments with a probability *r*, which characterizes the risk level of collective failure. Accordingly, the payoffs of cooperators and defectors in a group of size *N* with *j*_*C*_ cooperators and *N* − *j*_*C*_ defectors can be summarized as

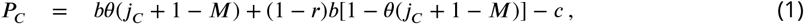

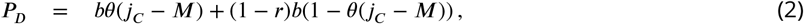

where *θ* (*x*) is the Heaviside function, that is, *θ* (*x*) = 0 if *x <* 0, being one otherwise.

To analyze the evolutionary dynamics of strategies in an infinite population, we use replicator equations to describe the time evolution of cooperation (***Taylor and Jonker, 1978***; ***Schuster and Sigmund, 1983***). Accordingly, we have

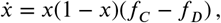

where *x* denotes the frequency of cooperators in the population, while *f*_*C*_ and *f*_*D*_ respectively denote the average payoffs of cooperators and defectors, which can be calculated as

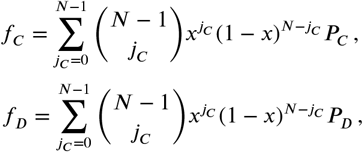

where *P*_*C*_ and *P*_*D*_ are shown in Eqs. (1) and (2). After some calculations, the difference between the average payoffs of cooperators and defectors can be written as

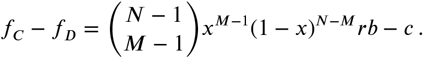

In the above replicator equation, we describe a game-theoretic interaction involving the risk of collective failure, which is a positive constant in previous works (***Santos and Pacheco, 2011***; ***Chen et al., 2012a***). Here, we are focusing on a dynamical system where there is feedback between strategic behaviors and risk. In particular, the impact of strategies on the risk level is channeled through a function *U* (*x, r*), which depends on both key variables. Then by using the general form of the feedback, the coevolutionary dynamics can be written as

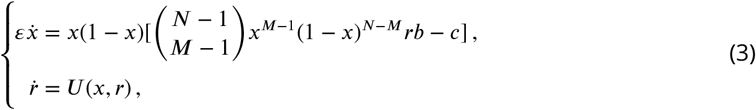

where *ε* denotes the relative speed of strategy update dynamics (***Weitz et al., 2016***), such that when 0 < *ε* ≪ 1 the strategies evolve significantly faster than the change in the risk level. In the following, we consider both linear and nonlinear forms of feedback describing the effect of strategy distribution on the evolution of risk.

### Linear effect of strategy on risk

In the first case, we assume that the effect of strategies on the risk level takes a linear form, which is the most common form that can be used to describe the characteristic attributes between key variables. Just to illustrate it by a specific example, the probability of influenza infection among individuals who have not been vaccinated decreases linearly with the increase of vaccine coverage (***Vardavas et al., 2007***). Here we consider that the value of risk decreases linearly with the increase of cooperation level. Furthermore, by following the work of Weitz et al. (***Weitz et al., 2016***), we can write the dynamical equation of risk as

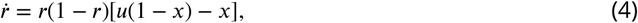

where *u*(1 − *x*) − *x* denotes the increase of risk by the defection level at rate *u* and the decrease by the fraction of cooperators at relative rate one. Then the dynamical system is described by the following equation

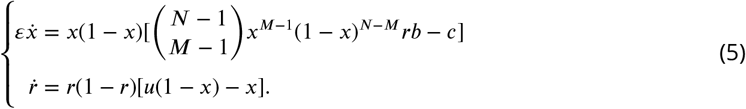

### Exponential effect of strategy on risk

To complete our study we also apply nonlinear form of feedback function. The most plausible choice is when the risk level depends exponentially on the population state. To be more specific, we consider that the risk will decrease when the frequency of cooperators in the population exceeds a certain threshold value *T*. Otherwise, the risk level will increase. Such scenario is suitable for describing climate change and the spread of infectious diseases, in which the risk can increase sharply, such as the occurrence of extreme weather (***Eckstein et al., 2021***) or a sudden outbreak of an epidemic in a region (***Yang and Shaman, 2022***). Here, we use the sigmoid function to describe the effect of strategy on the risk state (***Boza and Számadó, 2010***; ***Chen et al., 2012b***; ***Couto et al., 2020***), which can be written as

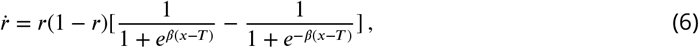

where *β* characterizes the steepness of the function and *r*(1 − *r*) ensures that the risk state remains in the [0, 1] domain. For convenience, we introduce the variable *ξ* = *x* − *T* and the function 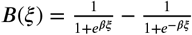. Thus we have 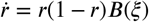. When *β* = 0, we know that *B*(*ξ*) = 0. In this situation, strategies have no effect on the risk level. For *β* = +∞, the function *B*(*ξ*) becomes steplike so that the risk will decrease only if the frequency of cooperators in the group exceeds the threshold *T*. Otherwise, the risk level remains high. To study the consequence of a proper feedback effect we apply a finite *β >* 0 value. In Figure 2 we illustrate how *B*(*ξ*) varies with *ξ* for four different values of *β*.

**Figure 2.**
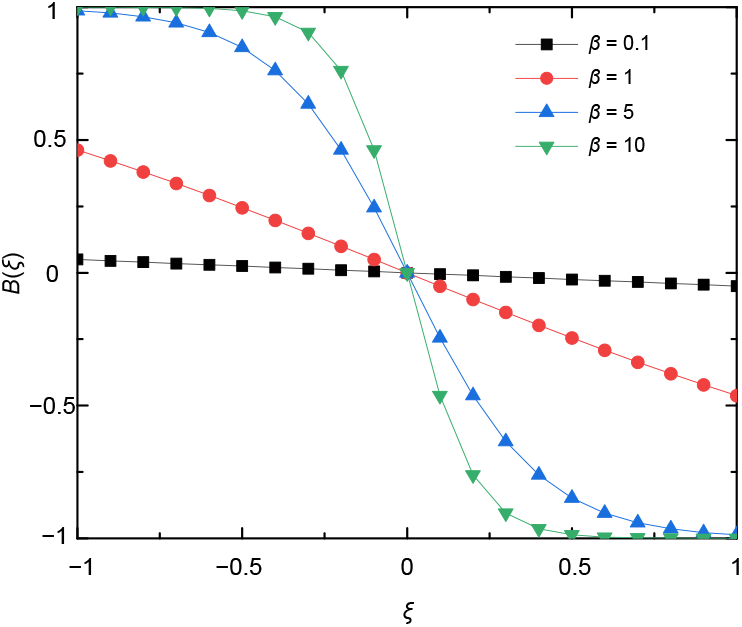
Feedback equation *B*(*ξ*) varies with *ξ* for different values of *β*. The parameter *β* determines the steepness of the curves. When the value of *β* is small, the *B*(*ξ*) function is almost constant or decays linearly by increasing *ξ*. For larger *β* values, the shape of *B*(*ξ*) approaches a step-like form. In this parameter area the risk level depends sensitively on whether the group cooperation exceeds the threshold *T* value or not.

Accordingly, the feedback-evolving dynamical system where the effect of strategies on the risk state is expressed by the exponential form can be written as

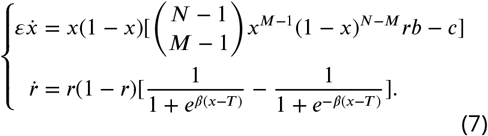

In the following section, we respectively investigate the coevolutionary dynamics of strategy and risk when considering linear and exponential feedback froms. We note that the details of theoretical analysis can be found in the Appendix.

## Results

### System I: coevolutionary dynamics with linear feedback

We first consider the case of linear feedback. More precisely, we assume that the risk value of collective failure will decrease linearly with the increase of cooperation and increase linearly with the increase of defection level. The resulting dynamical system is presented in Eq. (5). After some calculations, we find that this equation system has at most seven fixed points, which are (0, 0), (0,1), (1,0), (1,1), 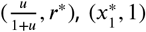 and 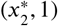, where 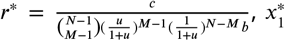 and 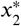 are the real roots of the equation 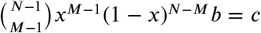. We further perform theoretical analysis for these equilibrium points, as provided in Appendix 1. In order to describe the stable states of System I for the complete parameter regions, we present a schematic plot in the parameter space 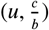, as shown in Fig. 3. We use different colors to distinguish the evolutionary outcomes for specific pairs of key parameters. In the following, we discuss the representative results in detail.

**Figure 3.**
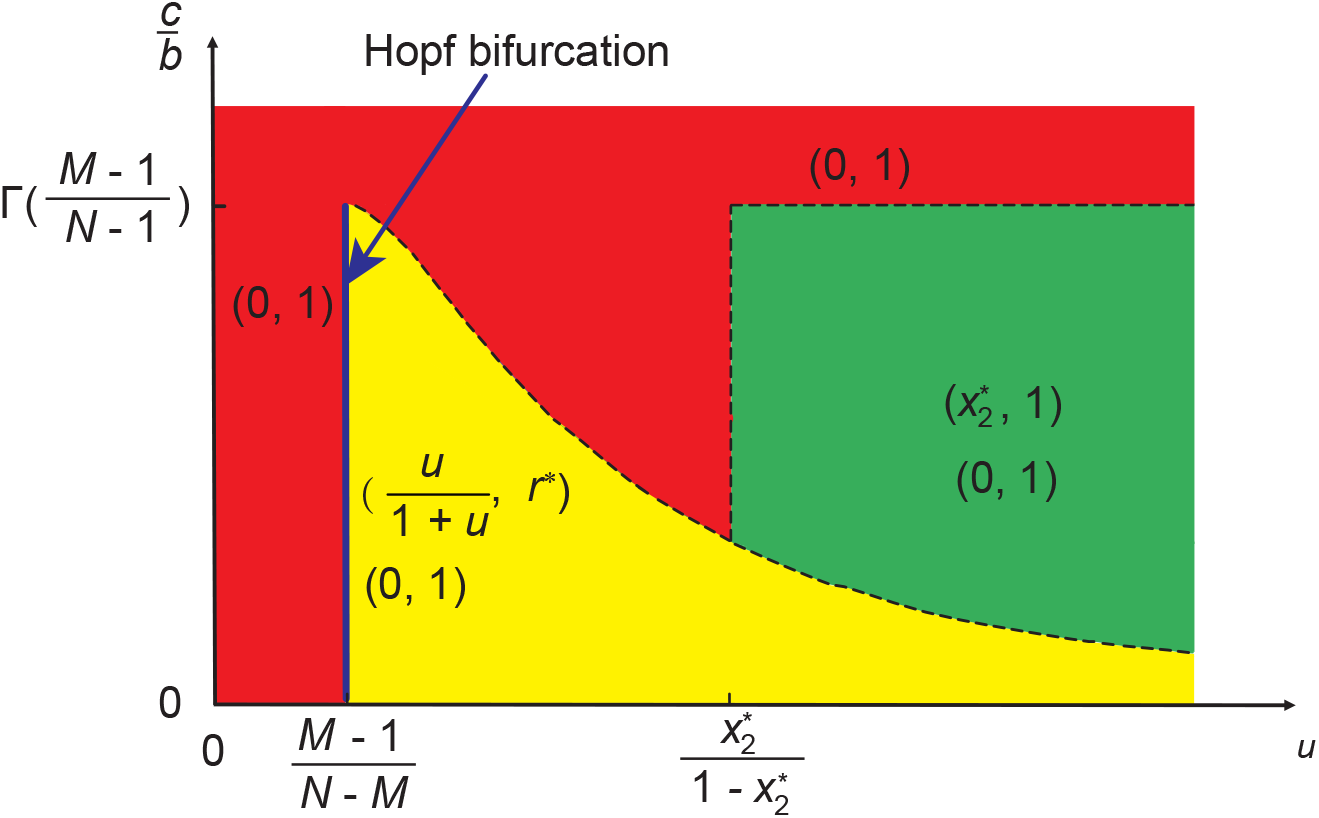
Representative plot of stable evolutionary outcomes in System I when linear strategy feedback on risk level is assumed. Different colors are used to distinguish the stability of different equilibrium points in the parameter space 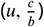. The blue line indicates that the system undergoes a Hopf bifurcation at 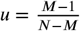. Here 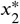 is the real root of the equation 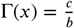 where 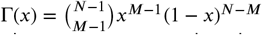, and 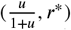 is the interior fixed point where 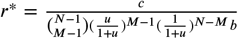. The dashed curve represents that the value of 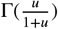 changes with *u* when 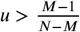. The horizontal dashed line represents that 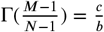 when 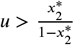. The vertical dashed line represents that 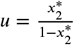 when 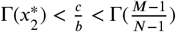.

#### System I has an interior equilibrium point

When 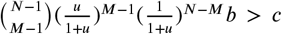, we know that our coevolutionary system has an interior fixed point. According to its stability, we can distinguish three sub-cases here. Namely, when 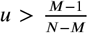, then the existing interior fixed point is stable. Besides, since 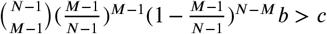, there exist seven fixed points in the system, namely, (0, 0), (0, 1), (1, 0), (1, 1), 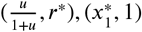, and 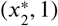. Here only (0, 1) and 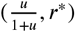 are stable (marked by yellow area in Fig. 3). Besides, we provide numerical examples to illustrate the above theoretical analysis (see the top row of Fig. 4). We find that bistable dynamics can appear, that is, depending on the initial conditions the system will evolve to one of two stable equilibria: here (0, 1) which means high-risk without cooperators, or the interior fixed point suggests that cooperation can be maintained at a high level when the value of risk exceeds an intermediate value. Furthermore, we note that the results are not affected qualitatively by the feedback speed (see the second column of Fig. 1 in Appendix 1).

**Figure 4.**
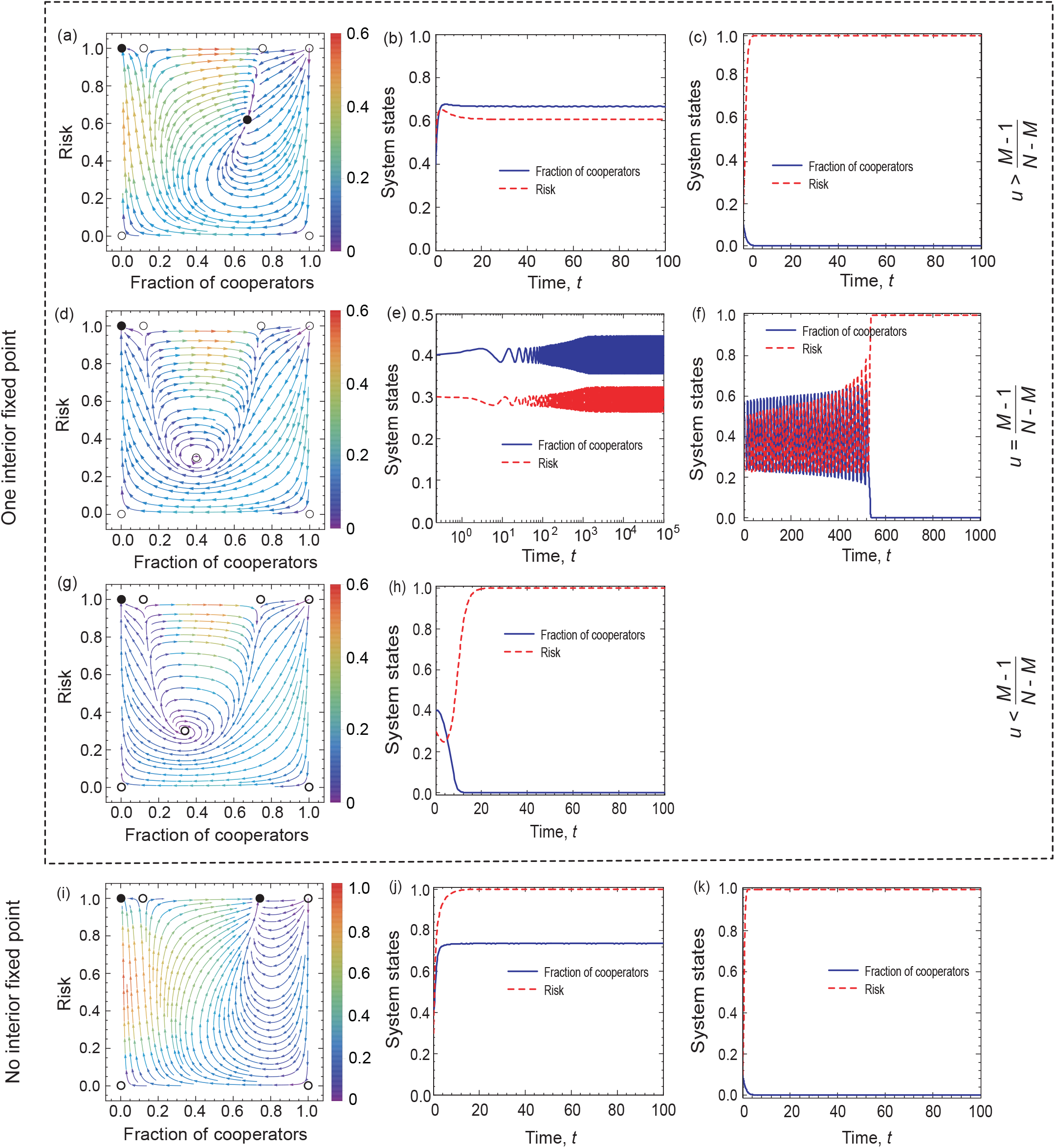
Coevolutionary dynamics on phase planes and temporal dynamics of System I when linear feedback is considered. Filled circles represent stable and open circles denote unstable fixed points. The arrows provide the most likely direction of evolution and the continuous color code depicts the speed of convergence in which red denotes the highest speed, while purple represents the lowest speed of transition. On the right-hand side, blue solid line and red dash line respectively denote the fraction of cooperation and the risk level, as indicated in the legend. The first three rows show the coevolutionary dynamics when 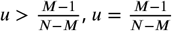, and 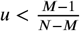, respectively. The bottom row shows coevolutionary dynamics when 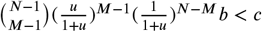. Parameters are *N* = 6, *c* = 0.1, *b* = 1, *μ* = 2, *ε* = 0.1, *M* = 3 in panel (a). The initial conditions are (*x, r*) = (0.4, 0.3) in panel (b) and (*x, r*)= (0.1, 0.1) in panel (c). *N* = 6, *c* = 0.1, *b* = 1, 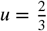 ε = 0.1, *M* = 3 in panel (d). The initial conditions are (*x, r*) = (0.4, 0.3) in panel (e) and (*x, r*) = (0.4, 0.5) in panel (f). *N* = 6, *c* = 0.1, *b* = 1, *μ* = 0.5, *ε* = 0.1,*M* = 3 in panel (g). The initial conditions are (*x, r*) = (0.4, 0.3) in panel (h). *N* = 6, *c* = 0.1, *b* = 1, *μ* = 4, *ε* = 0.1,*M* = 3 in panel (i). The initial conditions are (*x, r*) = (0.4, 0.3) in panel (j) and (*x, r*) = (0.1, 0.1) in panel (k).

If the enhancement rate of risk caused by defection drops to a certain threshold, namely 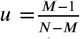, a Hopf bifurcation takes place, which is supercritical (marked by blue line in Fig. 3). In this situation, System I has all seven fixed points. As analyzed in Appendix 1, only (0, 1) is stable. Furthermore, we provide numerical examples to illustrate our theoretical analysis (see the second row of Fig. 4). We find that the system is bi-stable: depending on the initial fractions of cooperators and risk, the system can evolve either to a high-risk state without cooperation or to a limit cycle where the frequencies of cooperation and risk show periodic oscillations. In addition, we find that the key observations do not change qualitative by increasing the feedback speed (see the second column of Fig. 1 of Appendix 1).

When the enhancement rate of risk caused by defection is weak and meets 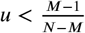 condition, then the interior fixed point is unstable. Besides, since 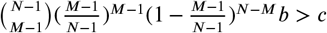, there exist all seven fixed points. According to the theoretical analysis presented in Appendix 1, only (0, 1) fixed point is stable. In the third row of Fig. 4, we present some representative numerical examples. They show that all trajectories in the state space terminate at the fixed point (0, 1), which is consistent with our theoretical results. This means that no individual chooses to contribute to the common pool, leading to the failure of collective action, and finally, all individuals inevitably lose all their endowments.

#### System I has no interior equilibrium point

The alternative case is when there is no interior fixed point, namely, 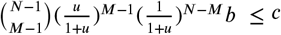. In this situation, when 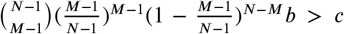, System I has six fixed points, which are (0, 0), (0, 1), (1, 0), (1, 1), 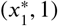, and 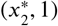, respectively. The theoretical analysis, presented in Appendix 1, shows that (0, 0), (1, 0), (1, 1), and 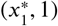 are unstable, (0, 1) is stable, and 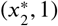 is stable for 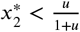 (shown by green area in Fig. 3). In the bottom row of Fig. 4, we provide some numerical examples to illustrate our theoretical results. The phase plane dynamics show that most trajectories in phase space converge to the stable equilibrium point 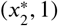, which suggests that driven by the high risk of future loss, most individuals will contribute to the common pool. Besides, the remaining trajectories in the phase space will converge to the fixed point (0, 1), which means a complete failure when all individuals lose all remaining endowments.

Furthermore, we prove that the fixed point 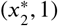 is unstable when 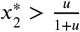 in Appendix 1. For the special case of 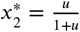, we find that one eigenvalue of the Jacobian matrix at 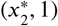 is zero and the other one is negative. We provide the stability analysis of this fixed point by using the center manifold theorem (***Khalil, 1996***). When 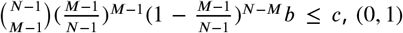 is the only stable equilibrium point of the System I.

### System II: coevolutionary dynamics with exponential feedback

In this section, we consider the case of exponential feedback. Here, there are at most seven equilibrium points of the replicator equation (7). Namely, (*x, y*) = (0, 0), (0, 1), (1, 0), (1, 1), 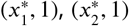, and 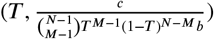, in which 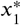 and 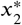 satisfy the equation 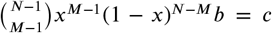 and 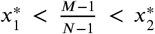 (***Santos and Pacheco, 2011***). For convenience, we set 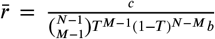. Here the first six equilibria are boundary fixed points, and the last one is an interior fixed point. In Appendix 2, we analyze the stability of these equilibria under four different parameter ranges by evaluating the sign of the eigenvalues of the Jacobian (***Khalil, 1996***). The basins of each solution in parameter space 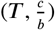 are shown in Fig. 5. In the following, we will discuss the evolutionary outcomes depending on whether System II has an interior equilibrium point.

**Figure 5.**
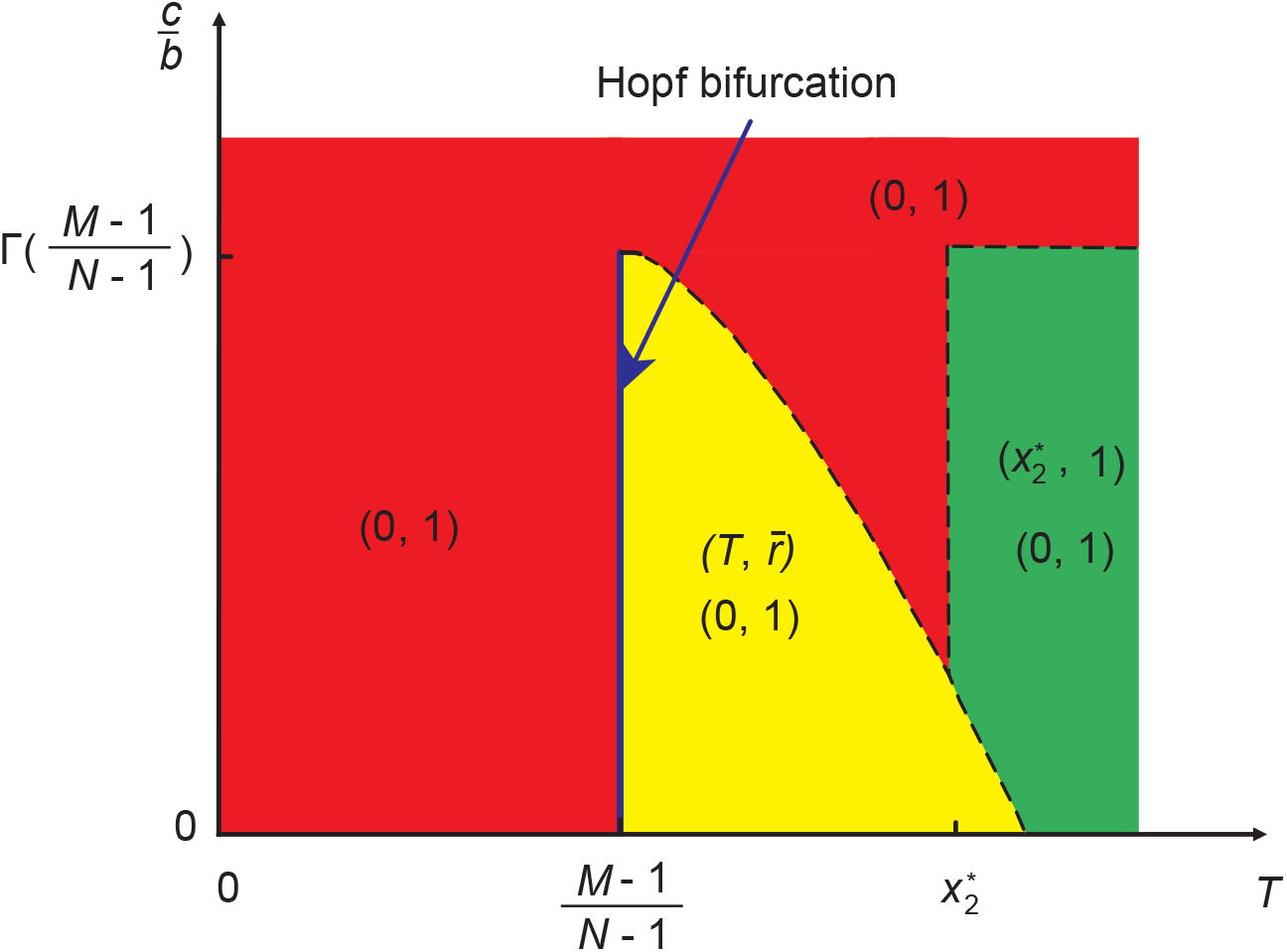
A representative diagram about stable solutions of System II when strategy feedback on risk level is exponential. We use different colors to distinguish the stability of equilibrium points in the parameter space 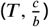. The blue line indicates that the system undergoes a Hopf bifurcation at 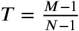. Here 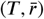 is the interior fixed point where 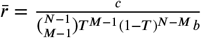. The dashed curve represents that the value of Γ (*T*) changes with *T* when 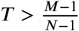. The horizontal dashed line represents that 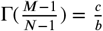 when 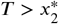. The vertical dashed line represents that 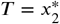 when 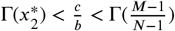.

#### System II has an interior equilibrium point

In this case 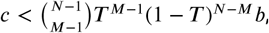, there are three typical dynamic behaviors for the evolution of cooperation and risk according to the stability conditions of the interior equilibrium point (for details, see Appendix 2).

When 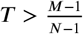, the interior fixed point is stable. Besides, since 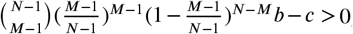, there exist two boundary fixed points, which are 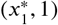 and 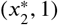. Thus the system has seven fixed points, which are (0, 0), (0, 1), (1, 0), (1, 1), 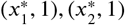, and 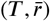). From the Jacobian matrices, we can conclude that the fixed points (0, 0), (0, 1), (1, 0), (1, 1), 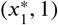, and 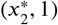 are unstable, while (0, 1) and 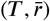 are stable. The latter case is shown in the top row of Fig. 6, where we plot the phase plane and temporal dynamics of the system. It suggests that there is a stable interior fixed point, and most trajectories in phase space converge to this nontrivial solution. Accordingly, the system can evolve into a state where the risk is kept at a low level and almost half of the individuals contribute to the common pool. The remaining trajectories in the phase space will converge to the alternative destination in which the risk level becomes particularly high and cooperators disappear. We note that these qualitative results are robust for different feedback speeds (see the first column of Fig. 1 in Appendix 2).

**Figure 6.**
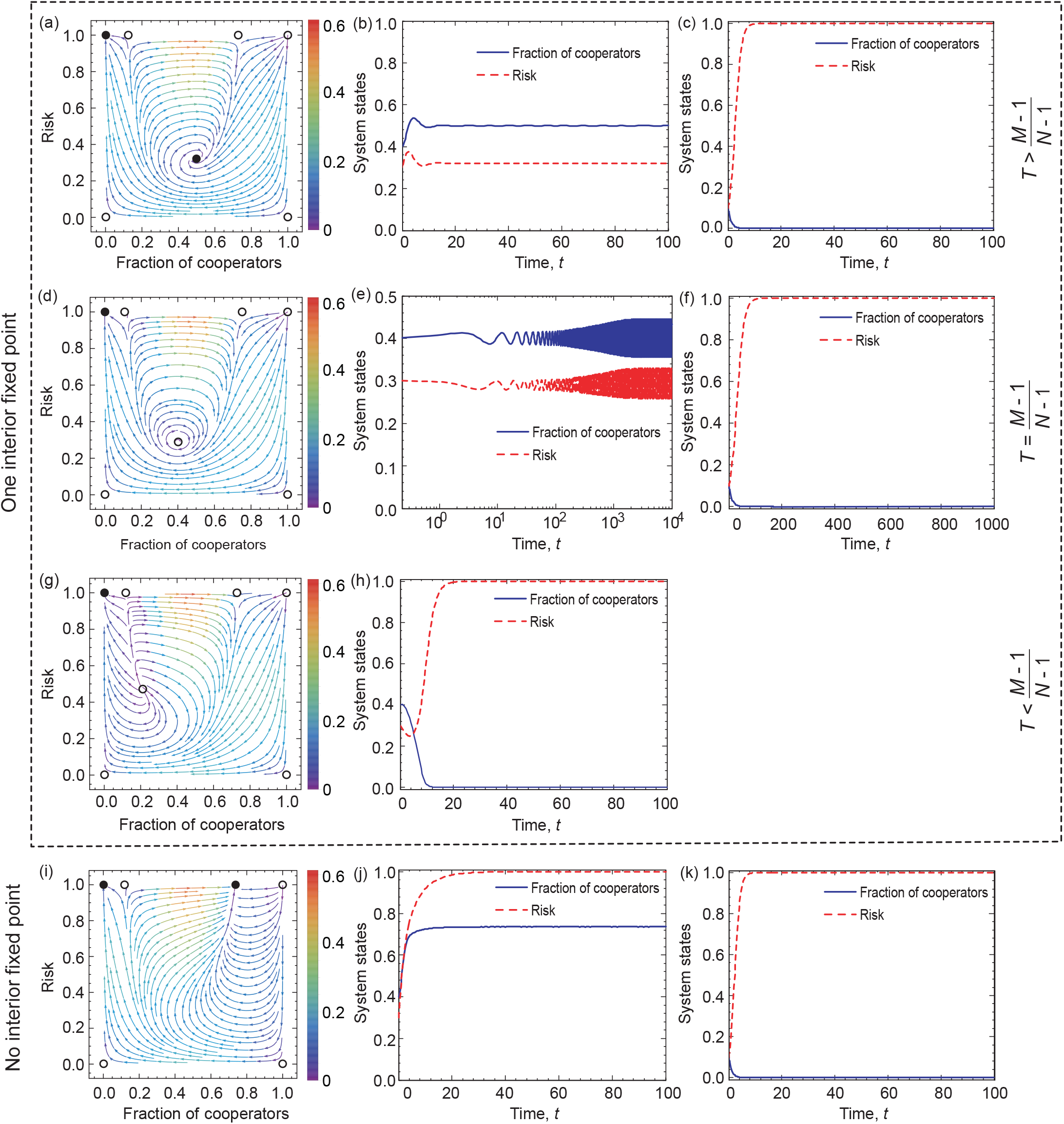
Coevolutionary dynamics on phase planes and temporal dynamics of System II when exponential feedback is assumed. Filled circles represent stable and open circles denote unstable fixed points. The arrows provide the most likely direction of evolution and the continuous color code depicts the speed of convergence in which red denotes the highest speed, while purple represents the lowest speed of transition. Blue solid line and red dash line respectively denote the fraction of cooperation and the risk level, as indicated in the legend. The first three rows show the coevolutionary dynamics when 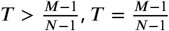, and 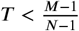, respectively. The bottom row shows the case when 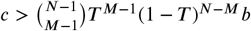. Parameters are *N* = 6, *c* = 0.1, *b* = 1, *T* = 0.5, *β* = 5, *ε* = 0.1,*M* = 3 in panel (a). The initial conditions are (*x, r*) = (0.4, 0.3) in panel (b) and (*x, r*) = (0.1, 0.1) in panel (c). *N* = 6, *c* = 0.1, *b* = 1, *T* = 0.4, *β* = 5, *ε* = 0.1,*M* = 3 in panel (d). The initial conditions are (*x, r*) = (0.4, 0.3) in panel (e) and (*x, r*) = (0.4, 0.5) in panel (f). *N* = 6, *c* = 0.1, *b* = 1, *T* = 0.2, *ε* = 5, *ε* = 0.1, *M* = 3 in panel (g). The initial conditions are (*x, r*) = (0.4, 0.3) in panel (h). *N* = 6, *c* = 0.1, *b* = 1, *T* = 0.8, *ε* = 0.1, *M* = 3 in panel (i). The initial conditions are (*x, r*) = (0.4, 0.3) in panel (j) and (*x, r*) = (0.1, 0.1) in panel (k).

When 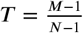, the eigenvalues of Jacobian matrix at the interior fixed point are a purely imaginary conjugate pair. Then, according to the Hopf bifurcation theorem (***Kuznetsov et al., 1998***; ***Guckenheimer and Holmes, 2013***), the system undergoes a Hopf bifurcation at 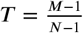 and a limitcycle encircling around interior equilibrium emerges. By calculating the first Lyapunov coefficient, we can evaluate that the limit cycle is stable (see Appendix 2). Besides, there exist two boundary fixed points, 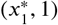 and 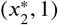, because 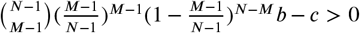. Thus the system has all seven fixed points. As we discuss in Appendix 2, only the fixed point (0, 1) is stable. A representative numerical example is shown in the second row of Fig. 6, which is conceptually similar to those we observed for System I. More precisely, the population either converges toward a limit cycle in the interior space, or arrives to the undesired (0, 1) point where there are no cooperators, but just high risk. Last, similar to previous cases, we observe that the feedback speed does not affect the behavior qualitatively, as illustrated in the second column of Fig. 1 of Appendix 2.

The interior fixed point is unstable when 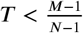. Besides, there are two boundary fixed points, 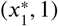 and 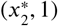, because 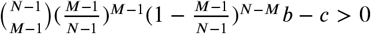. In this situation, the system has all seven fixed points. Theoretical analysis, presented in Appendix 2, confirms that only (0, 1) is stable. This is illustrated in the third row of Fig. 6 where all trajectories terminate in the mentioned point, signaling that the tragedy of the commons state is inevitable.

#### System II has no interior equilibrium point

When 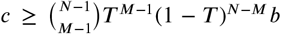, there is no interior fixed point in System II. In this case, when 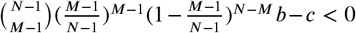, there are four equilibrium points, namely, (0, 0), (0, 1), (1, 0), (1, 1) where (0, 1) is stable. When 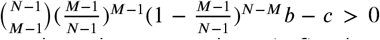, there exist two boundary fixed points, 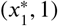 and 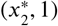. Altogether, the system has six fixed points, which are (0, 0), (0, 1), (1, 0), (1, 1), 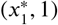, and 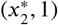. As we discuss in Appendix 2, the fixed points (0, 0), (1, 0), (1, 1), 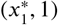 are unstable, while (0, 1) is stable. In the special case of 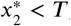, the fixed point 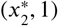 becomes stable. In this exotic state there is a significant cooperation at a high risk level. A representative numerical illustration is shown in the bottom row of Fig. 6, signaling the importance of the initial conditions, because the trajectories converge either to the fixed point (0, 1) or to 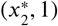.

## Discussion

Human behavior and the natural environment are inextricably linked. Motivated by this fact, rapidly growing research efforts have recognized the importance of developing a new comprehensive framework to study the coupled human-environment ecosystem (***Stern, 1993***; ***Liu et al., 2007***; ***Farahbakhsh et al., 2022***). Starting from the powerful concept of coevolutionary game theory, several works focus on depicting the reciprocal interactions and feedback between human behaviors and natural environment - both the impact of human behaviors on nature and the effects of environment on human behaviors (***Weitz et al., 2016***; ***Chen and Szolnoki, 2018***; ***Tilman et al., 2020***). Along this research line, we have developed a feedback-evolving game framework to study the coevolutionary dynamics of strategies and environment based on collective-risk dilemmas. Here, the environmental state is no longer a symbol of resource abundance, but depicts the risk level of collective failure. More precisely, we assume that the frequencies of strategies directly affect the risk level and reversely, the change of risk state stimulates individual behavioral decision-making. Importantly, we have explored both linear and highly nonlinear feedback mechanisms which characterize the link between the main system variables.

In particular, we have incorporated the strategies-risk feedback mechanism into replicator dynamics and explored the possible consequences of coevolutionary dynamics. We have shown that sustainable cooperation level can be reached in the population in two different ways. First, the coevolutionary dynamics can converge to a fixed point. This fixed point can be in the interior, indicating that the frequency of cooperators and the level of risk can be respectively stabilized at a certain level, or at the boundary, indicating high-level cooperation can be maintained even at a significantly high-risk environment. Second, the system has a stable limit cycle where persistent oscillations in strategy and risk state can appear. In addition, we have found that the above described evolutionary outcomes do not depend significantly on the character of feedback mechanism how strategy change affects on risk level. No matter it is linear or nonlinear, what really counts is the existence of the proper feedback. Importantly, we have theoretically identified those conditions which are responsible for the final dynamical outcomes. Interestingly, it is worth emphasizing that the relative evolutionary speed of strategy and risk level does not alter the behavior qualitatively, hence underlines the robustness of our observations.

Previous theoretical studies have revealed that the coevolutionary game models describing the complex interactions between collective actions and environment can produce periodic oscillation dynamics (***Weitz et al., 2016***; ***Tilman et al., 2020***). Although our feedback-evolving game model can also produce persistent oscillations, there are some differences. In particular, we have theoretically proved that Hopf bifurcation can take place and a stable limit cycle can appear in the system, which is different from the heteroclinic cycle dynamics reported by Weitz et al. (***Weitz et al., 2016***). Besides, we have found that the existence of a limit cycle does not depend on the speed of coupling, whereas Tilman et al. (***Tilman et al., 2020***) reported the opposite conclusion. Furthermore, we observe that a small amplitude oscillation is more conducive to maintaining the stability of the system than a large magnitude oscillation because a higher risk will make it easier for all individuals to lose all their endowments.

The reciprocal feedback process, though many types have not been well characterized, occurs at all levels of our life (***Liu et al., 2007***; ***Ezenwa et al., 2016***; ***Obradovich and Rahwan, 2019***). Consequently, they may play an indispensable role in maintaining the stability of human society and the ecosystem. Mathematical modeling based on evolutionary game theory is a powerful tool for addressing social-ecological and human-environment interactions and analyzing the evolutionary dynamics of these coupled systems. The mathematical framework proposed in this paper considers two characteristic forms to describe the effect of strategy on risk, namely, linear and nonlinear (exponential) forms of feedback. Although these two forms can be equivalent under some limit conditions, there are essential differences. On one hand, linear relationship is a relatively simple way to describe the correlation mode of two factors, which is common in real society. For example, with the increase of protection awareness and vaccination proportion, the mortality rate of the epidemic decreased gradually (***Yang and Shaman, 2022***). Furthermore, linear feedback has been used to describe the interactions between actions of the population and environmental state (***Weitz et al., 2016***; ***Tilman et al., 2020***). However, linear link cannot fully describe the relationship between variables in real societies. For example, in recent years, extreme weather phenomena have occurred more frequently, with greater intensity and wider impact areas. Thus the feedback between human behaviors and environment may take on a more complex nonlinear form. In this work, we consider that the strategy of the population has an exponential effect on risk level, and such form can describe the phenomenon that risk will rise and fall sharply with the change of strategy frequency (Fig. 2). It is worth emphasizing that although we use different forms of feedback to describe the impact of strategies on risk, the evolutionary dynamics have not changed substantially which highlights the prime importance of the feedback mechanism independently of its actual form.

Our feedback-evolving game model reveals that the coupled strategy and environment system will produce a variety of representative dynamical behaviors. We find that the undesired equilibrium point (0, 1) in our feedback system is always evolutionarily stable, which does not depend on whether the effect of strategy on risk is linear or exponential. Such evolutionary outcome means that all individuals are unwilling to contribute to achieving the collective goal, which leads to the failure of collective action, and all individuals inevitably lose their remaining endowments. In real-world scenarios, such as climate change (***Milinski et al., 2008***) and the spread of infectious diseases (***Cronk and Aktipis, 2021***; ***Chen and Fu, 2022***), once the whole society is in such a state, it is undoubtedly disastrous for the public. Therefore, how to adjust and control the system to deviate from this state is particularly important for policymakers.

Finally, it is worth emphasizing that the feedback loop operates over time. In this situation, the change of risk state or strategy frequency may lead to the change of other factors, such as collective target, which provides an opportunity for the emergence of new feedback loops. Thus, multiple types of feedback loops are possible in a single coupled system. Such multiple feedback loops have been confirmed in the coupling system of animal behavior and disease ecology (***Ezenwa et al., 2016***). Therefore, a promising expansion of our current model could be to consider the multiple feedback loops.

## Acknowledgments

This research was supported by the National Natural Science Foundation of China (Grants Nos. 61976048 and 62036002) and the Fundamental Research Funds of the Central Universities of China. L.L. acknowledges the support from Special Project of Scientific and Technological Innovation (Grant Nos. 2452022107 and 2452022012).

## Author contributions

Linjie Liu, Conceptualization, Formal analysis, Validation, Investigation, Methodology, Writing - review and editing; Xiaojie Chen, Conceptualization, Formal analysis, Supervision, Writing - review and editing; Attila Szolnoki, Formal analysis, Writing - review and editing

## Data Availability Statement Code availability

The Mathematica (Wolfram Mathematica 11.1) and Matlab (Matlab R2014a) source codes used to generate Figures 4 and 6 is available at the Dryad (https://doi.org/10.5061/dryad.wdbrv15rz).

## Appendix 1

We first study the case where the strategy of the population has a linear effect on the risk level. Then the dynamical system can be written as

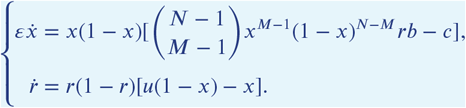

This equation system has at most seven fixed points, which are (0, 0), (0, 1), (1, 0), (1, 1), 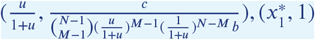, and 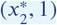, where 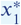 and 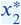 are the real roots of the equation 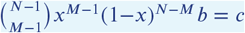. For convenience, we introduce the abbreviation 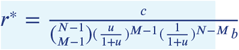 and 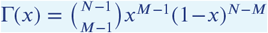. In the following, we analyze the stability of these equilibrium points.

(1) When 0 < *r*^*^ *<* 1, namely, 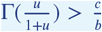, the system has an interior fixed point. Accordingly, the Jacobian for the interior fixed point is

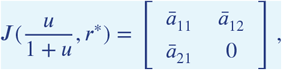

where 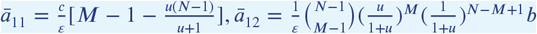, and 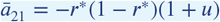.

(i) When 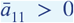, namely, 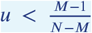, the existing interior fixed point is unstable. Since 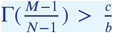, we can know that the two boundary fixed points 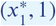 and 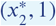 exist. Thus, the system has seven fixed points in the parameter space, namely, (0, 0), (0, 1), (1, 0), (1, 1), 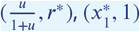, and 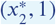. The Jacobian matrices of these equilibrium points are respectively given as follows.

For (*x, r*) = (0, 0), the Jacobian is

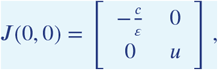

thus the fixed equilibrium is unstable.

For (*x, r*) = (0, 1), the Jacobian is

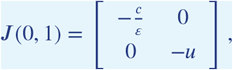

thus the fixed equilibrium is stable.

For (*x, r*) = (1, 0), the Jacobian is

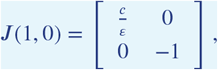

thus the fixed equilibrium is unstable.

For (*x, r*) = (1, 1), the Jacobian is

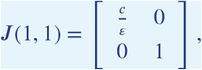

thus the fixed equilibrium is unstable.

For 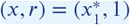, the Jacobian is

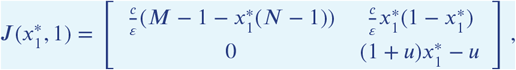

thus the fixed equilibrium is unstable since 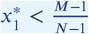.

For 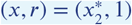, the Jacobian is

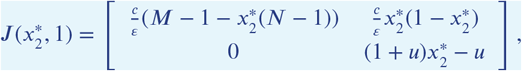

because 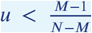 and 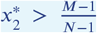, then 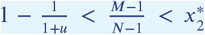. Thus this fixed equilibrium is unstable.

(ii) When 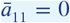, namely, 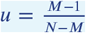, the trace and determinant of the Jacobian matrix at the interior equilibrium point are respectively given by

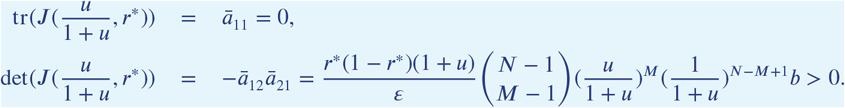

The eigenvalues of the Jacobian matrix can be calculated

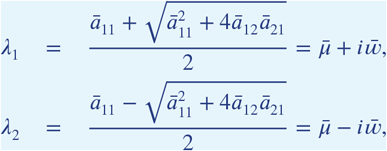

where 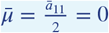 and 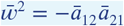.

Accordingly, we know that the eigenvalues satisfy the following conditions

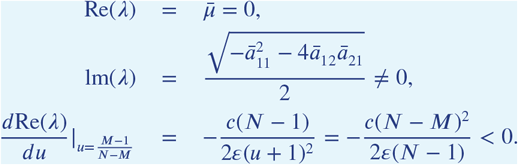

The first two conditions imply that the eigenvalues of Jacobian matrix at 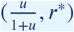 has a pair of pure imaginary roots. The third condition means that the pair of complex-conjugate eigenvalues crosses the imaginary axis with nonzero speed. According to Hopf bifurcation theorem (***Kuznetsov et al., 1998***), we know that a Hopf bifurcation takes place at 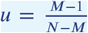. In order to determine the stability of the existing limit cycle from Hopf bifurcation, we need to calculate the first Lyapunov coefficient. We denote that 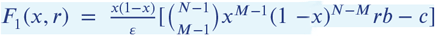 and 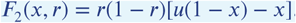.

Let *q, p* ∈ *𝒞*^2^ respectively denote the eigenvectors of the Jacobian matrix *J* (*T, r*^*^) and its transpose,

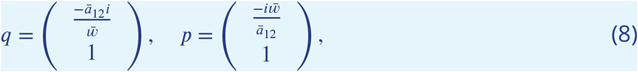

which satisfy

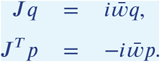

To achieve the necessary normalization 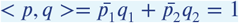, we can take

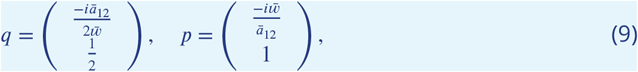

According to Ref. (***Kuznetsov et al., 1998***), we construct the complex-valued function

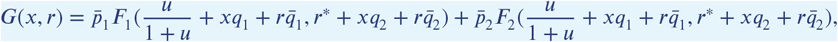

where *p, q* are given above, to evaluate its formal partial derivatives with respect to *x, r* at (*T, r*^*^), obtaining *g*_20_ = *G*_*xx*_, *g*_11_ = *G*_*xr*_, and *g*_21_ = *G*_*xxr*_. After some calculations, we can get the first Lyapunov coefficient

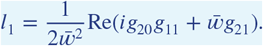

Specifically, when *l*_1_ *<* 0, a unique and stable limit cycle bifurcates from the equilibrium appears, while when *l*_1_ *>* 0, the Hopf bifurcation is subcritical such that an unstable limit cycle will be generated. Due to the complexity of the system, it is difficult to conduct bifurcation analysis collectively. Here we conduct a numerical analysis to investigate the stability of the existing limit cycle when the model parameters are consistent with Fig. 4(d). By using the algorithm in Ref. (***Kuznetsov et al., 1998***), we can get *l*_1_ = −1.407166124 × 10^−8^ *<* 0, which implies that the Hopf bifurcation is supercritical.

Besides, since 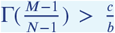, we can state that the two boundary fixed points 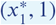 and 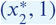 exist. Thus the system has seven equilibrium points, which are (0, 0), (0, 1), (1, 0), (1, 1), 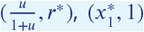, and 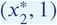, respectively. Accordingly to the sign of the eigenvalues of the Jacobian matrices, we know that only (0, 1) is stable.

(iii) When 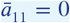, namely, 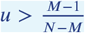, the trace and determinant of the Jacobian matrix at the interior equilibrium point are respectively given by

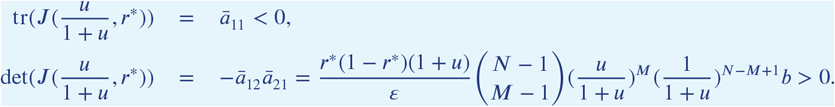

Thus the interior fixed point is stable. Besides, since 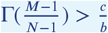, two boundary fixed points, 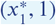 and 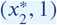, exist. Thus there are seven fixed points in the system, which are (0, 0), (0, 1), (1, 0), (1, 1), 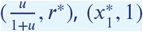, and 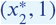, respectively. Here, the fixed points (0, 1) and 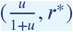 are stable, while others are unstable.

(2) When *r*^*^ 1, namely, 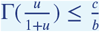, the system has no interior equilibrium point. In this case, when 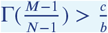, the system has six fixed points, which are (0, 0), (0, 1), (1, 0), (1, 1), 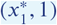, and 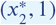, respectively. According to the sign of the largest eigenvalues of the Jacobian matrices, we can say that (0, 0), (1, 0), (1, 1), 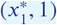 are unstable, while (0, 1) is stable. Particularly, when 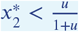, the fixed point 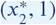 is stable, and it is unstable when 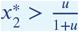. When 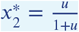, we know that one eigenvalue of the Jacobian matrix is zero and the other eigenvalue is negative. Then we study its stability by using the center manifold theorem (***Khalil, 1996***). For the fixed point 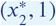, the Jacobian matrix can be written as

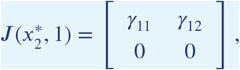

where 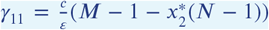 and 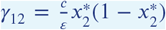. To do that, we take 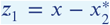 and *z*_2_ = *r* − 1, then the system can be rewritten as

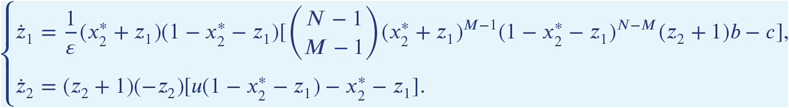

Let *Q* be a matrix whose columns are the eigenvectors of 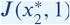, which can be written as

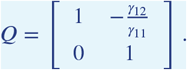

Then we have

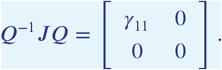

We further take [*η*_1_ *η*_2_]^*T*^ = *Q*^−1^[*z*_1_ *z*_2_], and then we have 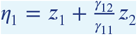 and *η*_2_ = *z*_2_. It leads to

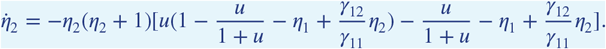

According to the center manifold theorem, we know that *η*_1_ = *h*(*η*_2_) is a center manifold. Then we start to try *h*(*η*_2_) = *0*(|*η*_2_|^2^), which yields the reduced system

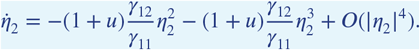

Since 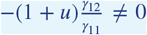, the fixed point *η*_2_ = 0 of the reduced system is unstable. Accordingly, the fixed point 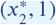 of the original system is unstable.

When 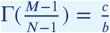, the system has five fixed points, which are (0, 0), (0, 1), (1, 0), (1, 1), and 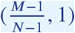, respectively. According to the sign of the eigenvalues in the Jacobian matrices, we can state that only (0, 1) is stable. When 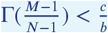, the system has four fixed points, namely (0, 0), (0, 1), (1, 0), and (1, 1). Here only (0, 1) is stable.

**Appendix 1 Figure 1.**
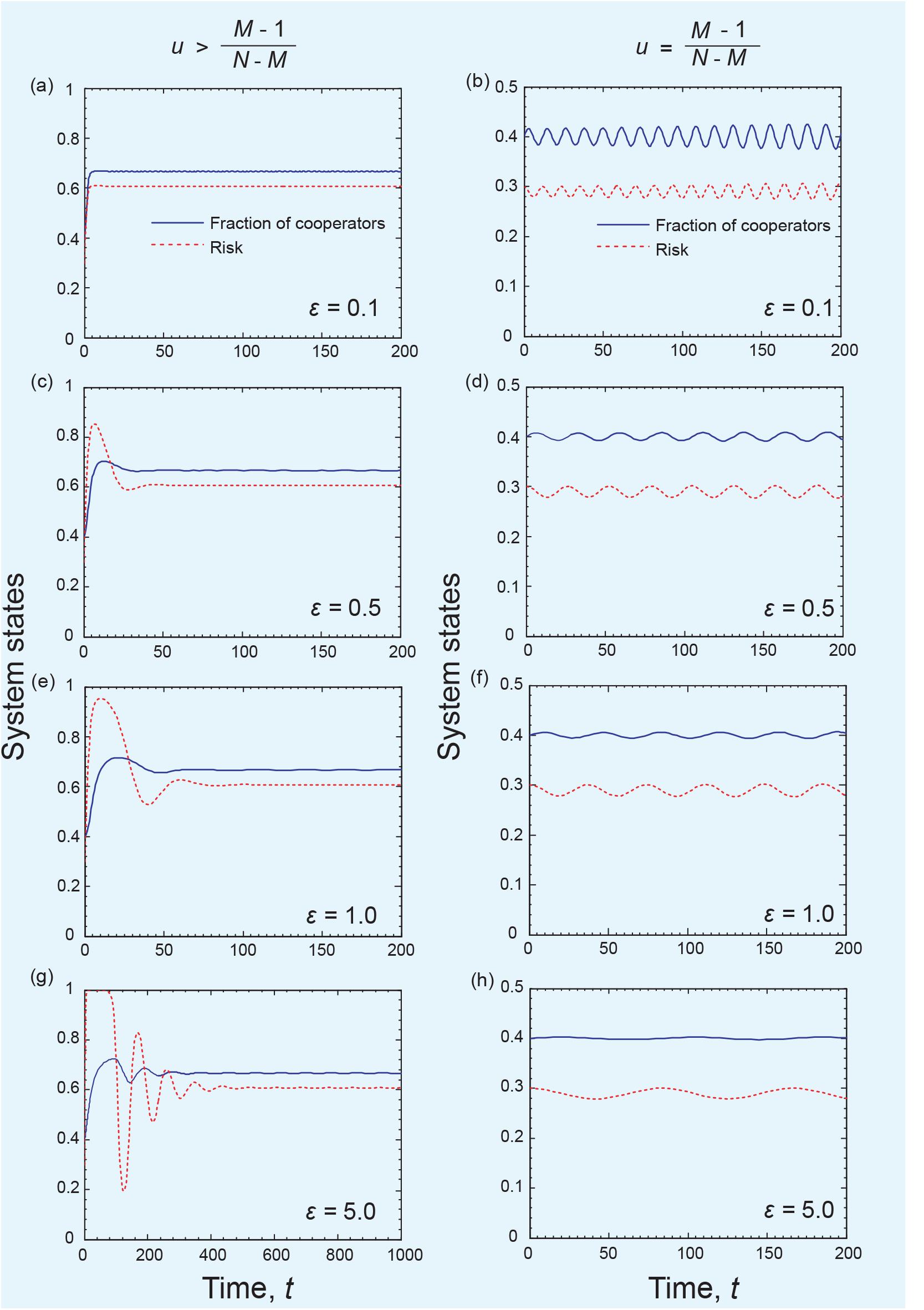
Coevolutionary dynamics of System I for different *ε* values when linear feedback effect of strategy on risk level is considered. Parameters are *N* = 6, *c* = 0.1, *b* = 1, and *M* = 3. *u* = 2 in left column and *u* = 2/3 in right column. The initial conditions are (*x, r*) = (0.4, 0.3).

## Appendix 2

System II with exponential feedback is described by

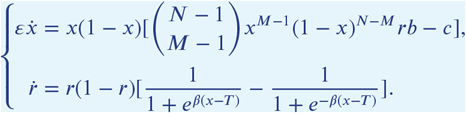

where the parameter *β >* 0 represents the steepness of the function.

This equation system has at most seven fixed points, which are (0, 0), (0, 1), (1, 0), (1, 1), 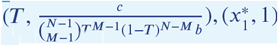, and 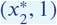, where 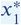 and 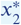 are the real roots of the equation 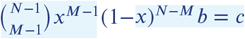 and 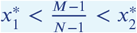. For simplicity, we introduce the abbreviation 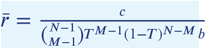 and 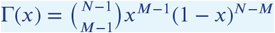. In the following, we study the stabilities of equilibria based on whether the system has an interior equilibrium point.

(1) When 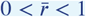, namely, 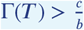, System II has an interior equilibrium point.

The Jacobian matrix evaluated at this equilibrium is

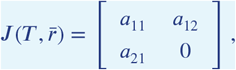

where 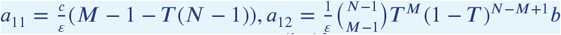, and 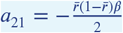. Notice that 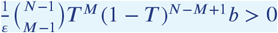 and 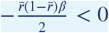, then the trace and determinant of the Jacobian matrix are respectively given by

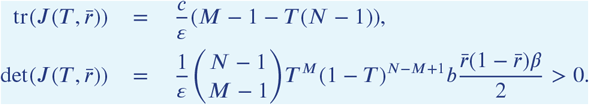

The eigenvalues of the Jacobian matrix can be calculated

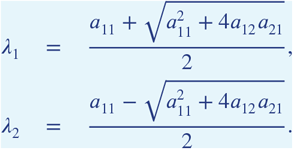

Here we set that 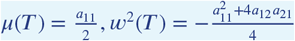, and 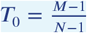

(i) When *a*_11_ *>* 0, namely, *T < T*_0_, the interior equilibrium point is unstable. Since 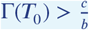, we can know that the two boundary fixed points 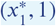 and 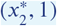 exist. Thus, the system has seven fixed points in the parameter space, namely, (0, 0), (0, 1), (1, 0), (1, 1), 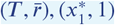, and 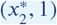.

For (*x, r*) = (0, 0), the Jacobian is

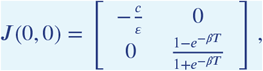

thus the fixed equilibrium is unstable.

For (*x, r*) = (0, 1), the Jacobian is

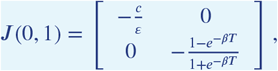

thus the equilibrium point is stable.

For (*x, r*) = (1, 0), the Jacobian is

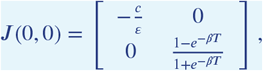

thus the fixed point is unstable.

For (*x, r*) = (1, 1), the Jacobian is

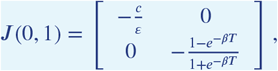

thus the fixed equilibrium is unstable.

For 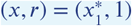, the Jacobian is

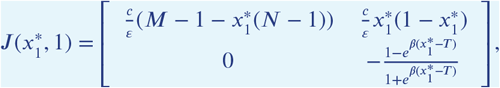

thus the fixed equilibrium is unstable since 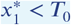.

For 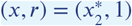, the Jacobian is

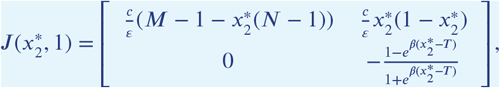

thus the fixed equilibrium is unstable since 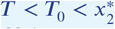.

(ii) When *a*_11_ = 0, namely, 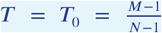, we have *μ*(*T*_0_) = 0. Moreover, 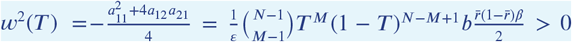. Therefore, the eigenvalues of the Jacobian matrix are a purely imaginary conjugate pair *λ*_1,2_(*T*_0_) = ± *iw*(*T*_0_). Considering that 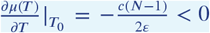, then we know that the system undergoes a Hopf bifurcation at *T* = *T*_0_ and there exists a limit cycle around the interior equilibrium. Accordingly, we can evaluate the direction of the limit cycle bifurcation by computing the first Lyapunov coefficient *l*_1_ of the system. Here we also conduct numerical calculations to investigate the stability of the existing limit cycle when the model parameters are consistent with Fig. 6(d). By using the algorithm in Ref. (***Kuznetsov et al., 1998***), we can get *l*_1_ = −1.876221498 × 10^−8^, which implies that the Hopf bifurcation is supercritical.

Besides, since 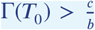, we know that there are seven equilibrium points in System II. They are (0, 0), (0, 1), (1, 0), (1, 1), 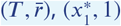, and 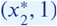. According to the sign of the eigen-values of the Jacobian matrices, only (0, 1) is stable.

(iii) When *a*_11_ *<* 0, namely, *T > T*_0_, the interior equilibrium point is stable. Besides, since 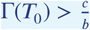, we find that there are seven fixed points in the system, which are (0, 0), (0, 1), (1, 0), (1, 1), 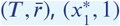, and 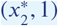, respectively. Here, the fixed points (0, 1) and 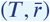 are stable, while others are unstable.

(2) When 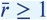, namely, 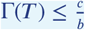, System II has no interior equilibrium point. In this case, when 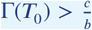, the system has six fixed points, which are (0, 0), (0, 1), (1, 0), (1, 1), 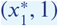, and 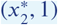, respectively. According to the sign of the largest eigenvalues of the Jacobian matrices, we can say that (0, 0), (1, 0), (1, 1), 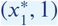 are unstable, while (0, 1) is stable. Particularly, when 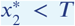, the fixed point 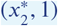 is stable, and it is unstable when 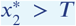. When 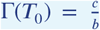, the system has five fixed points, which are (0, 0), (0, 1), (1, 0), (1, 1), and (*T*_0_, 1), respectively. According to the sign of the eigenvalues in the Jacobian matrices, we can see that only (0, 1) is stable. When 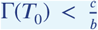, the system has four fixed points, namely (0, 0), (0, 1), (1, 0), and (1, 1). Here only (0, 1) is stable.

**Appendix 2 Figure 1.**
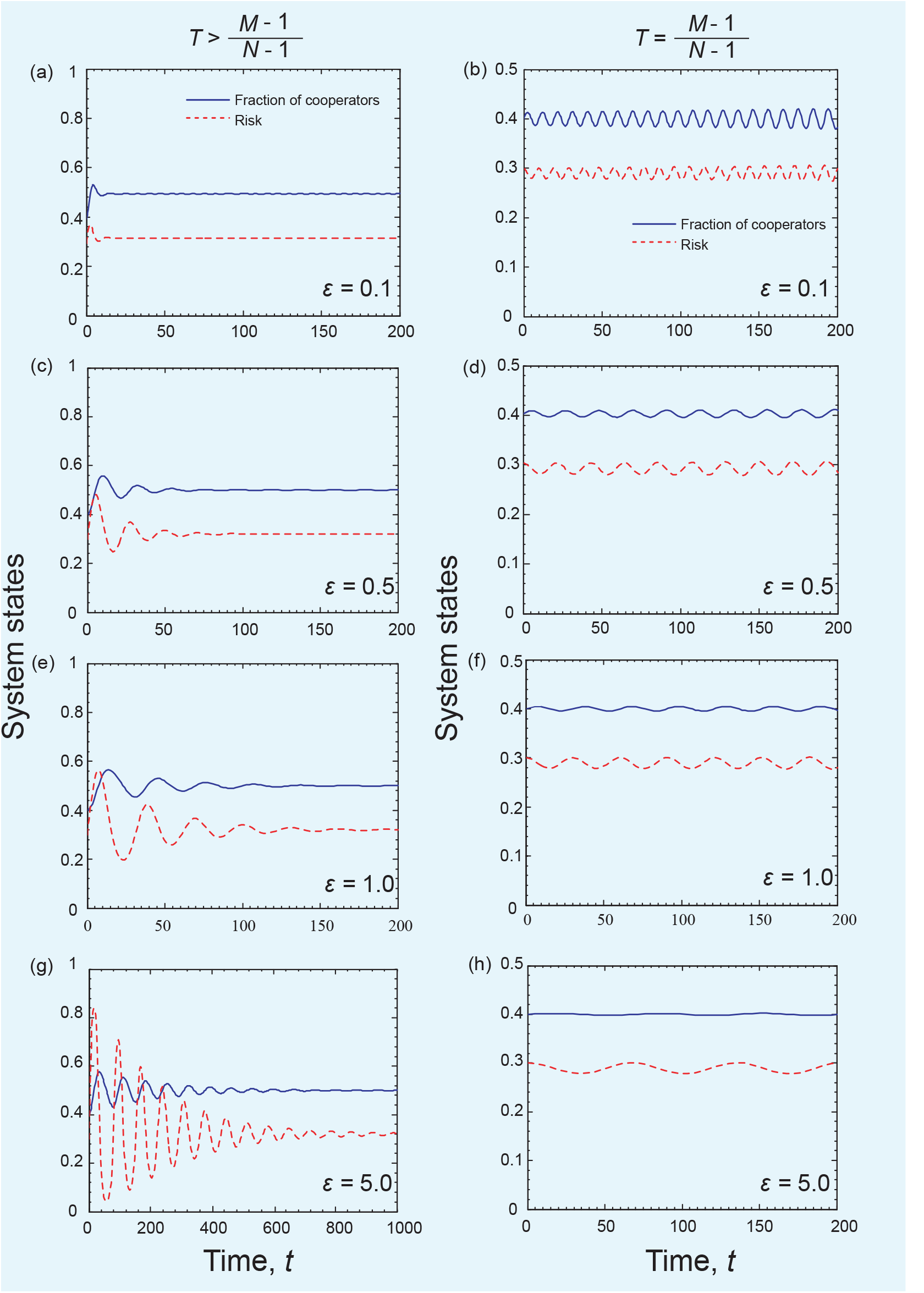
Coevolutionary dynamics of System II for different *ε* values when the strategy feedback on risk is exponential. Parameters are *N* = 6, *c* = 0.1, *b* = 1, *β* = 5, and *M* = 3. *T* = 0.5 in the left column and *T* = 0.4 in the right column. The initial condition is (*x, r*) = (0.4, 0.3).

